# Neuroligins mediate presynaptic maturation through brain-derived neurotrophic factor signaling

**DOI:** 10.1101/262246

**Authors:** Andoniya Petkova, Nina Gödecke, Martin Korte, Thomas Dresbach

**Author notes:** Corresponding author and Lead Contact: Thomas Dresbach, Tel. +49 (0)551/397004, Fax +49 (0)551/397043.

## Abstract

Maturation is a process that allows synapses to acquire full functionality; failures in synaptic maturation may contribute to neurodevelopmental disorders. Neuroligins are postsynaptic cell-adhesion molecules essential for synapse maturation, but the underlying pathways are incompletely understood. Here, we show that maturational increases in active zone stability and synaptic vesicle recycling rely on the joint action of Neuroligins and brain-derived neurotrophic factor (BDNF). Applying BDNF to neuronal cultures mimicked the maturation-promoting effects of overexpressing the Neuroligin isoforms NL1 or NL2. Inhibiting BDNF signaling reduced the effects of NL1 and NL2 on presynaptic maturation and of NL2 on synapse formation. Applying BDNF to NL1-knockout mouse cultures rescued defective presynaptic maturation, indicating that BDNF acts downstream of NL1 and can restore presynaptic maturation at late stages of network development. Our data introduce BDNF as a novel and essential component in a transsynaptic pathway linking Neuroligin-mediated cell adhesion, neurotrophin action and presynaptic maturation.

## Introduction

Synaptic maturation is a complex event that turns newly formed synapses into fully functional units. Maturation endows synapses with adequate levels of stability, strength and plasticity, and optimizes them for their specific roles in a certain network. Activity-dependent maturation shapes developing circuits, while impaired synapse maturation profoundly affects brain function, contributing to neurodevelopmental and neurodegenerative disorders(Baudouin et al. 2012; Zoghbi 2003; Bourgeron 2009).

Neuroligins are postsynaptic cell adhesion molecules that regulate the number and maturation of synapses. Both loss and gain of function of Neuroligins has been linked to the aetiology of autism spectrum disorders (ASD) (Südhof 2008) and Alzheimer’s disease (Betancur, Sakurai, and Buxbaum 2009; Gilman et al. 2011; Sindi, Tannenberg, and Dodd 2014), highlighting their importance for brain function. There are four principal isoforms in rodents (NL1, NL2, NL3 and NL4) and five in humans (NL1, NL2, NL3, NL4X, NL4Y), where NL1 is specific for excitatory synapses, while NL2 is primarily localized to inhibitory synapses. Neuroligins bridge the synaptic cleft by binding to their presynaptic receptors, the Neurexins. Increasing the levels of Neuroligins increases the number of synapses on the dendrites of the overexpressing neurons, while decreasing their levels reduces the number of synapses (Dean and Dresbach 2006; Südhof 2008). These effects require competition between cells, because they are only observed when Neuroligins are overexpressed or knocked down in a small number of neurons amidst unperturbed neurons (Chih, Engelman, and Scheiffele 2005; Kwon et al. 2012; Schnell et al. 2014; Shipman and Nicoll 2012; Futai et al. 2013).

In contrast, both global and cell-specific changes in the levels of Neuroligins affect synaptic maturation, indicating that promoting maturation represents a fundamental and essential function of Neuroligins. For example: global triple knockout of NLs 1, 2 and 3 perturbs synaptic maturation as evidenced by reduced GABA-receptor recruitment to inhibitory synapses. In addition, subsets of synapses are silent in these mice (Varoqueaux et al. 2006). Global knockout of NL1 reduces the strength of glutamatergic synapses by reducing NMDA receptor recruitment, and overexpression of NL1 in acute slices increases NMDA receptor recruitment, synaptic strength, and the stability of active excitatory synapses (Chubykin et al. 2007; Budreck et al. 2013). Likewise, overexpression of NL1 in vivo increases the size of dendritic spine heads and the recruitment of pre and postsynaptic proteins (Hoy et al. 2013; Dahlhaus et al. 2010).

How do Neuroligins exert their effects on synaptic maturation? We previously used cultured hippocampal neurons to determine in detail which aspects of synaptic maturation are regulated by Neuroligin 1 (Wittenmayer et al. 2009). Overexpressing NL1 increased NMDA receptor recruitment, presynaptic release probability, the number of presynaptic terminals with functional active zones (AZs), the number of recycling synaptic vesicles per terminal, and the stability of AZs in the absence of F-actin, all of which are hallmarks of maturation. These effects are reminiscent of the effects described for brain-derived neurotrophic factor (BDNF) in the context of neuronal development and plasticity. BDNF is a secreted transsynaptic signalling molecule of the neurotrophin family that binds to the TrkB (tropomyosine-related receptor kinase B) neurotrophin receptor and the pan-neurotrophin receptor p75^NTR^. In addition to its functions in promoting neuronal survival, neurite differentiation and axon guidance, BDNF mediates synaptic plasticity during postnatal development and in the mature nervous system (Park and Poo 2013; Zagrebelsky and Korte 2014). At the subcellular level, BDNF increases the number of synapses (Vicario-Abejón et al. 1998; W J Tyler and Pozzo-Miller 2001; Kellner et al. 2014), and enhances synaptic protein recruitment, vesicle recycling and release probability (Tartaglia et al. 2001; Collin et al. 2001; William J Tyler et al. 2006; Shinoda et al. 2014). We hypothesized that this potency of BDNF may mediate AZ maturation and interact with Neuroligin-based cell adhesion during network development.

Here, employing assays to test for structural and functional maturation, we found that applying BDNF to immature cultured neurons mimics the effects of overexpressing NL1 or NL2 on synapse number and maturation. Moreover, BDNF application rescued the maturational defects of the loss of NL1. Conversely, overexpressing NL1 or NL2 did not rescue defective maturation in BDNF-depleted cultures. In fact, the action of NL1 and NL2 on presynaptic maturation was severely impaired, suggesting that Neuroligins and BDNF act in concert to specifically mediate presynaptic maturation.

## Materials and methods

### Cell culture

Primary hippocampal neuron cultures were prepared from E19 Wistar rats and primary cortical neuron cultures were prepared from P0 NL1-WT and NL1-KO mice essentially as described previously (Wittenmayer et al. 2009). 55 000 cells/cm^2^ were plated on PEI-coated coverslips in 24-well plates. 12 hours after plating the original DMEM-medium containing 10% FCS, 100 µg/ml penicillin, 100 µg/ml streptomycin, and 1% L-glutamine was exchanged for Neurobasal medium with the same additives and B27 supplement.

Primary hippocampal mouse cultures were prepared from C57BL/6J-SV129 mice on E16.5. Immediately after having been taken out, the hippocampi were placed in ice-cold GBSS (Gey’s balanced salt solution; 137 mM NaCl, 5.55 mM D-Glucose, 4.89 mM KCl, 0.33 mM KH_2_PO_4_, 1.033 mM MgCl_2_*6H_2_O, 0.284 mM MgSO_4_*7H_2_O, 2.7 mM NaHCO_3_, 0.845 mM Na_2_HPO_4_, 1.98 mM CaCl_2_). After a 30-minute digestion in Trypsin/EDTA and 3–4 washes with serum medium (a 2% FCS solution in DMEM) the hippocampi were homogenized and then plated on poly-L-lysine coated coverslips at a 70 000 cells/cm^2^ density and kept in Neurobasal medium containing 0.25% Glutamine, 2% B27 supplement and 10% 10X N2 supplement. The cultures were incubated at 37°C and 5% CO_2_.

### Transfection

Neurons were transfected on day *in vitro* 2 (DIV2) or DIV4 using the calcium phosphate transfection method as described by Dresbach et al. 2003. In short, the original medium was first substituted with OptiMem medium; a mixture containing the plasmid, sterile distilled water, CaCl2, and the transfection buffer (274 mM NaCl, 10 mM KCl, 1.4 mM Na_2_HPO_4_, 15 mM Glucose, 42 mM HEPES, pH 7.06) was prepared and incubated for 20 minutes before being added to the cell medium. The cells were incubated for 60 minutes and then washed with Neurobasal medium. In the end, the original medium was restored. The following constructs were overexpressed: a farnesylated GFP; an mOrange construct; a Neuroligin-1 construct containing a GFP tag inserted in the intracellular domain (Dresbach et al. 2004) and a newly generated construct containing the NL1 sequence, an internal ribosomal entry site (IRES), and mOrange; a newly generated Neuroligin-2 construct with an mGFP tag attached to the C/terminus, i.e. to the intracellular domain, and a newly generated construct containing the NL2 sequence, and IRES, and mOrange. All constructs were inserted in a *pN1* vector.

### BDNF application

0.1 µg/ml BDNF (R&D Systems) was added to non-transfected neurons 22 hours before fixation. 1% BSA solved in sterile PBS (used to reconstitute BDNF) was added to the cell medium in the control condition.

### TrkB-Fc scavenger

TrkB-Fc (R&D Systems) dissolved in sterile PBS was added at a final concentration of 4 µg/ml to inhibit the TrkB receptor on DIV3 (12 hours after transfection).

### F-actin depolymerization

2.5 µM LatrunculinA (Sigma-Aldrich) dissolved in DMSO was applied to the medium 6 hours prior to fixation (Zhang and Benson 2001). To visualize F-actin TRITC-labeled phalloidin was used (1:6000).

### Synaptotagmin-1 antibody uptake

A monoclonal antibody directed against the lumenal domain of Synaptotagmin-1 (Synaptic Systems) was diluted (1:600) in depolarization buffer (64 mM NaCl, 70 mM KCl, 2 mM CaCl_2_, 1 mM MgCl_2_, 20 mM HEPES pH 7.4, 30 mM glucose) to reach a final dilution of 1:1200 in the cell medium. The neurons were then incubated for 5 minutes and subsequently washed three times with Neurobasal medium.

### Recombinant lentivirus production and infection

The production of the lentivirus was performed in HEK293T cells (5∙106 cells in 10 cm cell culture dish). The cells were co-transfected in serum free DMEM with 10 µg of the lentiviral expression vector (pUbc-Cre-IRES-GFP) in conjunction with the 7.5 µg of delta 8.9 vector and 5 µg of VSV-G using Polyethylenimine (PEI, Sigma Aldrich). The transfection solution was incubated for 30 minutes at RT and then drop-wise added to the cells. 4–6 h after the transfection culture medium was replaced by fresh DMEM containing 10% FCS. Lentivirus was harvested from the cell culture supernatant 48–72 h after transfection and immediately stored at -70°C. To excise the *bdnf* gene, primary hippocampal neurons from mice carrying two floxed bdnf alleles were transduced with the *cre*-recombinase expressing lentivirus. The lentivirus containing medium was added drop wise to the primary neurons in a 1:5 dilution. The transduction was performed 4 h after preparation of the primary hippocampal cultures.

### Immunofluorescence and microscopy

Neurons were fixed with 4% PFA in K^+^-free PBS for 20 minutes. Afterwards they were permeabelized using PBS with 0.3% Triton X-100, 2% BSA, 10% FCS, and 5% glucose.

Primary antibodies: mouse anti-Bassoon (Enzo Life Sciences) 1:1000; guinea pig anti-Synaptophysin (Sigma-Aldrich) 1:500; mouse anti Synaptotagmin-1 (Synaptic Systems) 1:200; mouse anti-BDNF (monoclonal antibody developed by Y.-A. Barde, obtained from the Developmental Studies Hybridoma BANK, created by the NICHD of the NIH and maintained at the University of Iowa, Department of Biology, Iowa City, IA 52242) 1:100. They were diluted in PBS with 0.3% Triton X-100, 2% BSA, 10% FCS, and 5% glucose.

Secondary antibodies: anti-mouse Alexa 647 1:1000; anti-mouse Cy3 1:500; anti-guinea pig Alexa 647 1:1000; anti-rabbit Alexa 647 1:1000. They were diluted in PBS with 0.3% Triton X-100, 2% BSA, and 5% glucose.

Microscope images were acquired using a CoolSNAP HQ2 CCD camera (Photometrics) attached to a Zeiss AxioObserver Z1 with a 40× magnification. Exposure time for the images with AZ and vesicle labeling was kept consistent for each experiment.

### Image analysis

900×900 px selections (the average size of 7 cell somas) of each image were made with Adobe Photoshop. Synaptic puncta were counted using OpenView (created by Noam E Ziv, Technion, Haifa). Synaptic puncta were selected automatically after setting a threshold. The threshold was set to 400, and then kept constant for all images. Dendritic length was measured in MetaMorph (Molecular Devices) and ImageJ. Synaptotagmin-1 intensity was measured using OpenView and ImageJ. The threshold was set in a way that all visible puncta in a neuron were selected and was kept constant throughout a set. Statistical analysis involved a two-tailed Student’s t-test and a two-way ANOVA and was performed using Prism (GraphPad Software).

## Results

### NL1, NL2 and BDNF induce structural presynaptic maturation

In cultured neurons, presynaptic maturation can be readily monitored using two time windows: before DIV7 (day in vitro 7), nerve terminals are largely immature, after DIV14 the majority of synapses have acquired a mature stage. During structural maturation, terminals become independent of F-actin, presumably by forming increasingly tight molecular complexes. The F-actin-disrupting drug Latrunculin-A (LatA) causes the loss of synaptic proteins from presynaptic terminals in young cultures but not in advanced cultures, providing a means to assay structural maturation (Zhang and Benson 2001; Wittenmayer et al. 2009). During functional maturation the total recycling pool of synaptic vesicles increases. Functional maturation can be readily determined using SV recycling assays (Mozhayeva et al. 2002; Wittenmayer et al. 2009). Taking advantage of these two time windows, we applied gain-of-function assays before DIV7 (asking what enhances and accelerates maturation) and loss-of-function assays after DIV14 (asking what prevents maturation).

Using these assays, we have previously shown that NL1 mediates structural and functional presynaptic maturation (Wittenmayer et al. 2009). To test if NL2 has similar potency, we transfected rat cultured neurons with NL1-GFP, NL2-GFP or membrane-targeted EGFP (farnesylated EGFP, EGFP-F) on DIV2. On DIV5 we fixed them with and without prior LatA treatment and immunostained for Bassoon to determine the number of AZs (Wittenmayer et al. 2009). Overexpressing NL1 or NL2 increased the number of Bassoon puncta per dendrite, indicating that both isoforms induce synapse formation (Fig. 1A). Virtually all Bassoon puncta formed on NL1 or NL2 expressing cells were LatA-resistant (Fig. 1B). Thus, both NL1 and NL2 accelerate structural AZ maturation. To test if BDNF promotes presynaptic terminal maturation, we applied BDNF to cultured neurons on DIV4, and determined the LatA resistance of Bassoon on DIV5. BDNF application increased the number of Bassoon puncta, suggesting that it caused the formation of AZs (Fig. 1C, D). Virtually all Bassoon puncta on BDNF-treated dendrites were LatA-resistant, reflecting advanced structural maturation. Thus, BDNF application and overexpression of NL1 or NL2 have similar effects in inducing structural maturation of AZs (Fig. 1).

**Figure 1:**
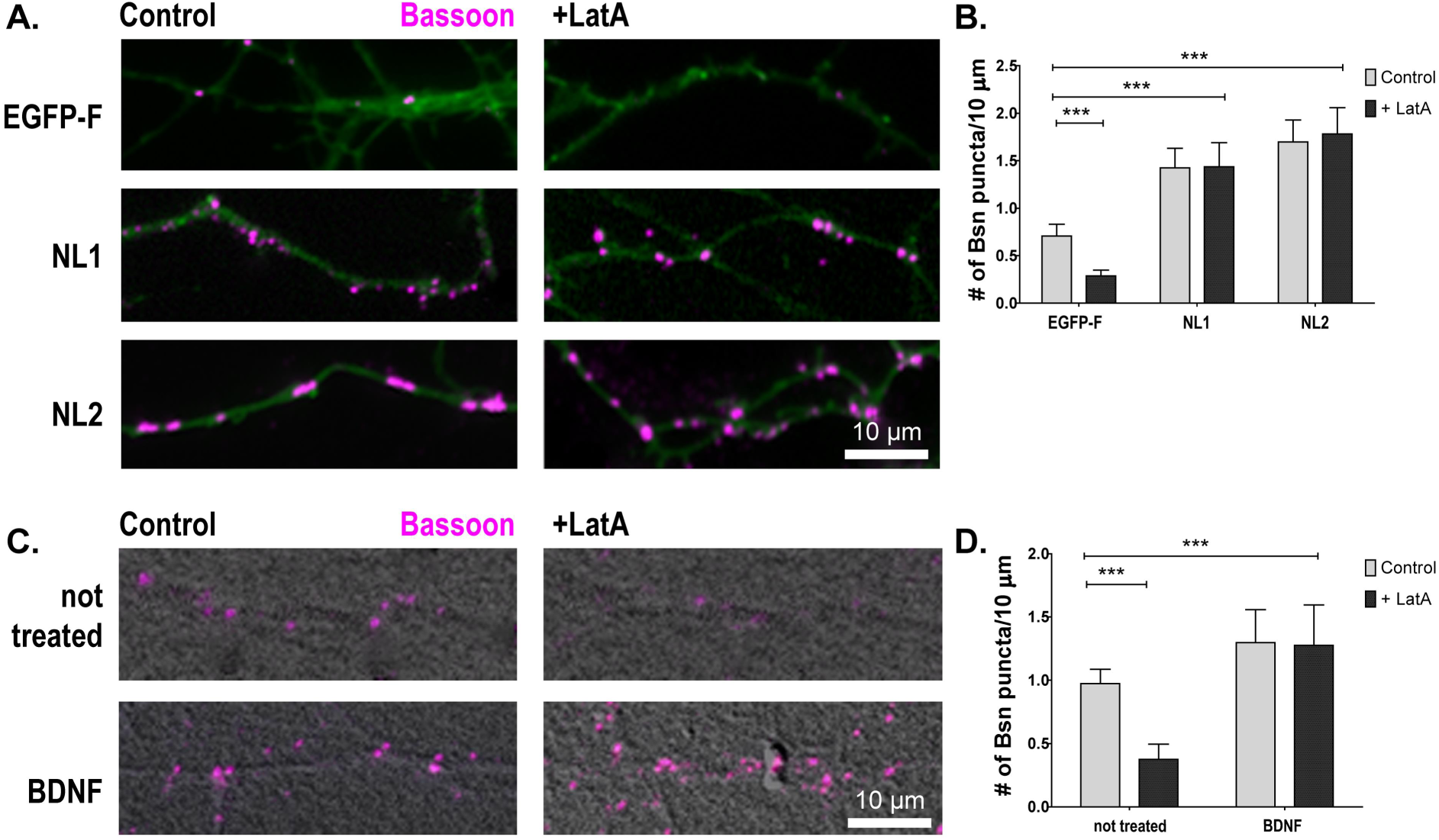
Induction of structural active zone maturation by NL1, NL2 and BDNF. (A) DIV5 cultured hippocampal neurons transfected with membrane targeted EGFP (farnesylated EGFP, EGFP-F), or GFP tagged NL1 or NL2 (green), with and without LatA treatment and immunostained for Bassoon (magenta). (B) Quantification of the number of Bassoon puncta per 10 µm dendrite for the conditions indicated in panel A. Virtually all Bassoon puncta formed on NL1 or NL2 expressing cells are LatA-resistant. (C) Cultured hippocampal neurons treated with BDNF or buffer (=not treated, NT) on DIV4, with and without LatA-treatment, and then fixed on DIV5, and immunostained for Bassoon (magenta). (D) Quantification of the number of Bassoon puncta per 10 µm dendrite for the conditions indicated in panel C. (Mean ± SD is shown; n = 3 experiments, 6 cells per experiment; ***p<0.001, paired t-test and two-way ANOVA (***p<0.001)). Scale bar is 10 µm.

### BDNF signaling is required for NL-induced structural and functional presynaptic maturation

To test if Neuroligins require BDNF signaling to promote AZ maturation, we overexpressed NL1 and NL2 in rat cultured neurons in the presence of TrkB-Fc, a soluble fragment of the BDNF receptor which is known to bind and ‛scavenge’ secreted BDNF, thus reducing the concentration of BDNF released into the media (Dean et al. 2009). TrkB-Fc reduced the number of Bassoon puncta on GFP-expressing cells, but did not block the action of NL1 in increasing puncta number. Remarkably, TrkB-Fc had distinct effects on the actions of NL1 and NL2, as it prevented the increase in puncta number induced by NL2 overexpression (Fig. 2A, B). Thus, BDNF appears to be required for synapse formation induced by NL2 but not for synapse formation induced by NL1. To test whether the Bassoon puncta formed under conditions of perturbed BDNF signaling represent structurally mature or immature AZs, we determined the LatA resistance of Bassoon in TrkB-Fc treated cultures. The small number of AZs formed by NL2 in the presence of TrkB-Fc was not further reduced by the LatA-treatment, suggesting that they are structurally mature. However, due to their small initial number we cannot exclude that we might miss a further reduction by LatA application. In contrast, TrkB-Fc clearly reduced the number of LatA-resistant Bassoon puncta on NL1-expressing cells (Fig. 2C, D), indicating that a significant proportion of AZs formed by NL1 remain structurally immature when BDNF signaling is perturbed.

**Figure 2:**
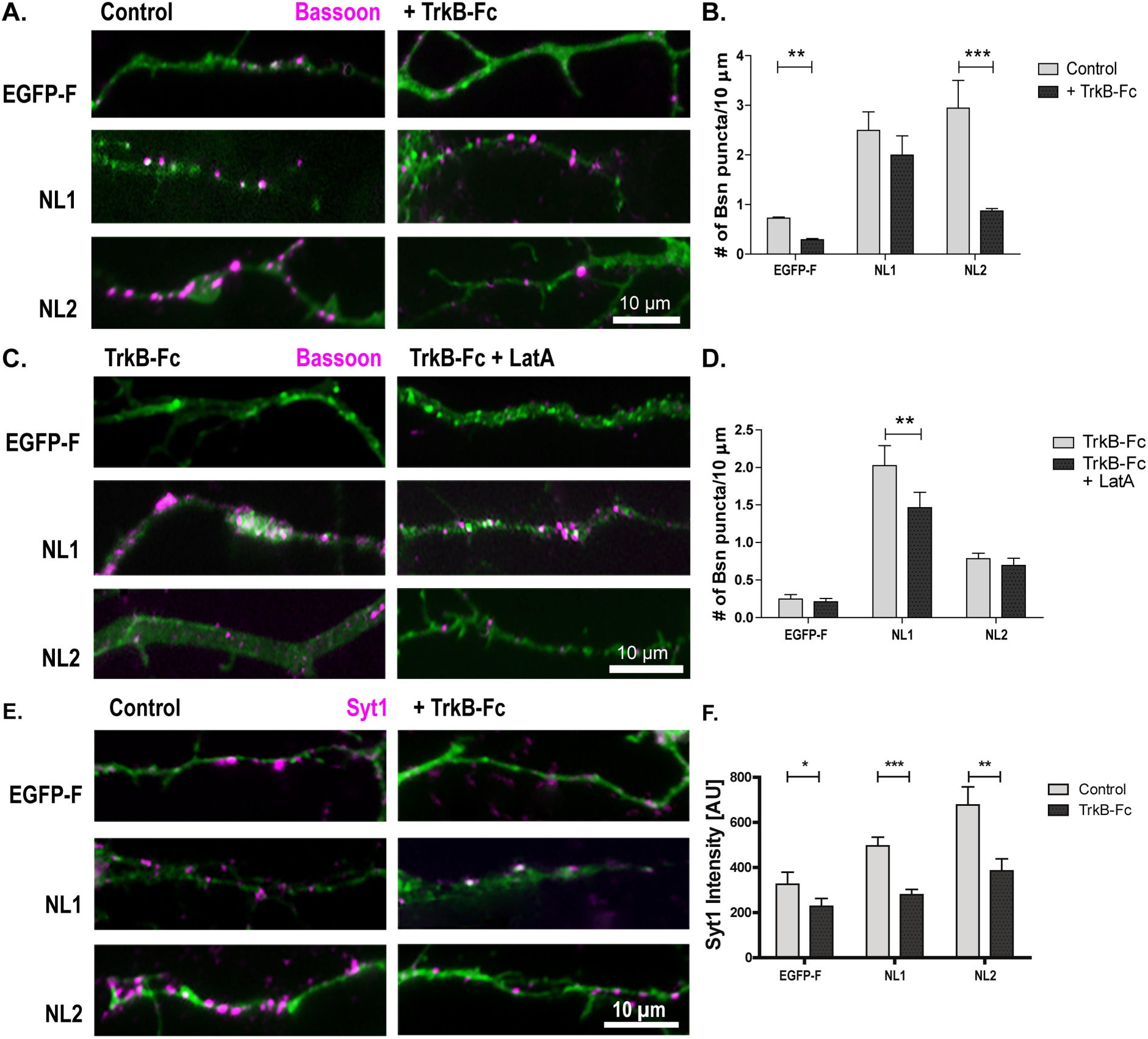
BDNF signaling is required for distinct aspects of NL1 and NL2 action. (A, B) DIV5 cultured hippocampal neurons transfected with farnesylated EGFP (EGFP-F), NL1-EGFP (NL1) or NL2-EGFP (NL2) (green) grown in the absence or presence of TrkB-Fc and immunostained for Bassoon (magenta), to determine the number of active zones. (C, D) TrkB-Fc treated cultures with and without LatA treatment immunostained for Bassoon, to determine the number of LatA-resistant active zones. (Mean ± SEM; n = 3 exp., 6 cells per exp.; **p<0.01, ***p<0.001 paired t-test). (E, F) BDNF signaling is required for NL mediated functional presynaptic maturation. Representative images and quantitation of cultured hippocampal neurons transfected on DIV2 with EGFP-F, NL1-EGFP (NL1), or NL2-EGFP (NL2) (green) and grown in the absence or presence of TrkB-Fc and stimulated in the presence of antibodies directed against the lumenal domain of Synaptotagmin-1 (Syt) to label recycling synaptic vesicles. After washing, the cells were fixed and immunolabeled with secondary antibodies against the Syt1-antibody (magenta). (Mean ± SEM; n = 3 exp., 6 cells per exp.; **p<0.01, ***p<0.001 paired t-test). Scale bar is 10 µm.

To test if BDNF signaling is required for functional maturation of NL-induced synapses, we assayed depolarization-driven uptake of anti-Synaptotagmin-1 antibodies. This assay exploits the fact that neurotransmitter release represents a cycle of exo- and endocytosis: upon exocytosis, the lumenal domains of synaptic vesicle transmembrane proteins become exposed at the plasma membrane surface of presynaptic nerve terminals. Bath applied antibodies that detect such lumenal domains are taken up into the nerve terminal during the endocytotic phase of neurotransmitter release and can be detected by fluorophore-coupled secondary antibodies. Thus, the fluorescence intensity is proportional to the number of antibodies taken up during the stimulus, and therefore reflects the number of vesicles undergoing the exo-endocytotic cycle. On DIV5, EGFP-F, NL1-EGFP, or NL2-EGFP expressing cultures were stimulated in the presence of antibodies directed against the lumenal domain of Synaptotagmin-1 (Syt1), to label recycling synaptic vesicles. After washing, the cells were fixed and immunolabeled with secondary antibodies against the Syt1-antibody. NL1 and NL2 increased the intensity of the Syt1 signal per terminal, indicating enhanced synaptic vesicle recycling in nerve terminals formed on NL1 and NL2 expressing cells (Fig. 2E, F). TrkB-Fc reduced Syt1-immunofluorescence in terminals in control cultures, and abolished the NL1- and NL2-induced increase in synaptic vesicle recycling.

### BDNF is required for presynaptic maturation

To further investigate how BDNF affects NL-induced presynaptic maturation, we tested DIV15 cultures from mice carrying two floxed *bdnf* alleles, where the BDNF gene was excised by transducing the cultures with *Cre*-recombinase-expressing lentivirus on the day of plating. In cultures from NL1 knockout mice presynaptic terminals fail to mature (Wittenmayer et al. 2009). If BDNF is part of the underlying pathway, BDNF depleted cultures should have impaired presynaptic maturation, too. To test this prediction, we determined the LatA resistance of AZs in DIV15 BDNF-depleted cultures. LatA application significantly reduced the number of Bassoon puncta in Cre-transduced cultures, but not in control cultures (Fig. 3A, B).

**Figure 3:**
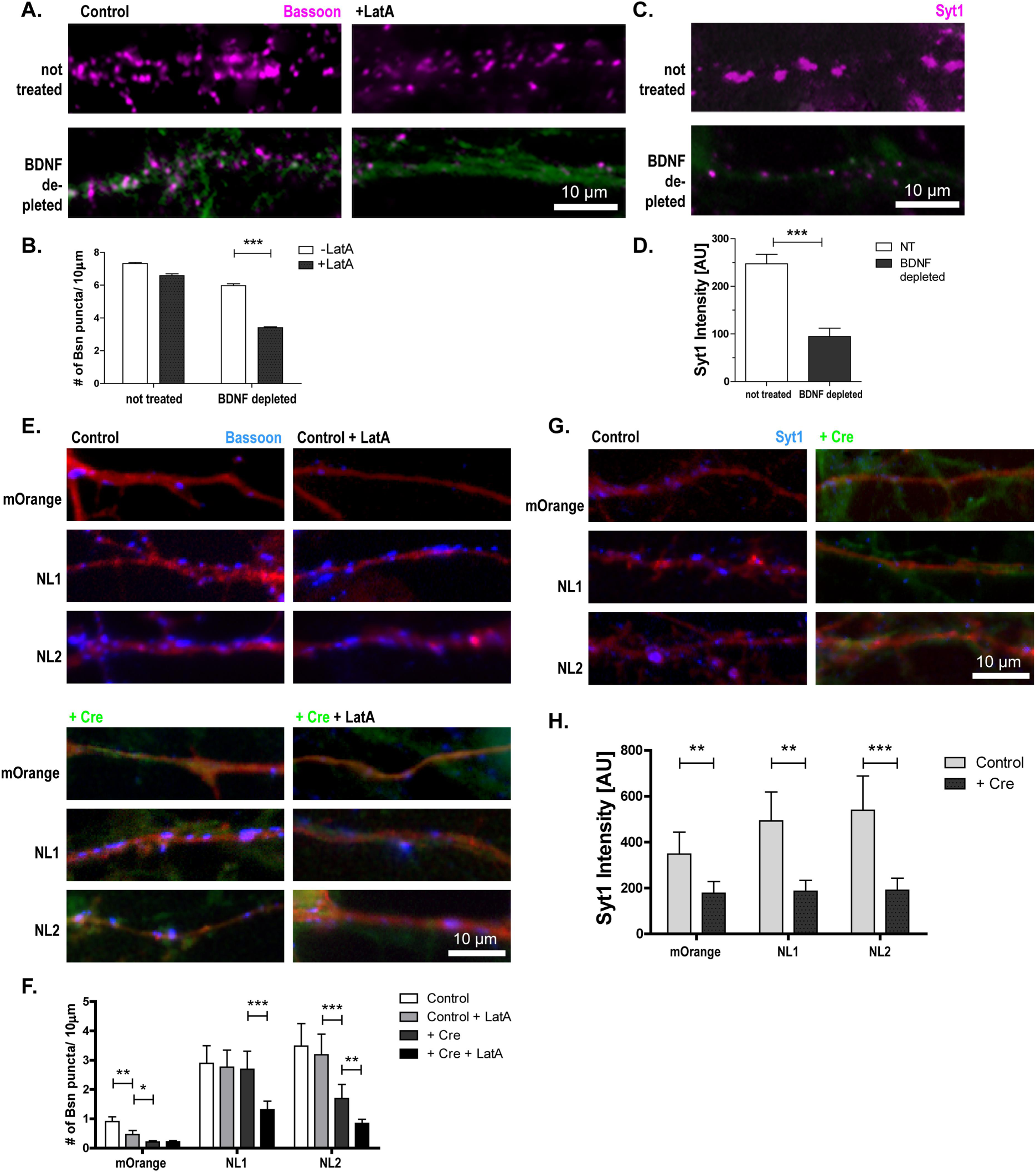
BDNF is required for endogenous and NL-induced presynaptic maturation. Cultures of BDNF^lox/lox^ hippocampal neurons with Cre-recombinase viral transduction (green) to reduce BDNF levels compared to non-transduced cultures. (A) Representative images of cultures fixed on DIV15 and immunostained for Bassoon (magenta) to determine the number of active zones following an 18-hour LatA treatment to depolymerize F-Actin. (B) Quantification of the number of Bassoon puncta per 10 µm dendrite for the conditions indicated in panel A. (Mean ± SEM; n = 3 exp., 6 cells per exp.; ***p<0.001, paired t-test). Scale bar is 10 µm. (C) Representative images of cultures of BDNF^lox/lox^ hippocampal neurons with Cre-recombinase viral transduction (green) to reduce BDNF levels compared to non-transduced cultures that were stimulated in the presence of antibodies directed against the lumenal domain of Synaptotagmin-1 (Syt1) to label recycling synaptic vesicles. After washing the cells were fixed and immunolabeled with secondary antibodies against the Syt1-antibody (magenta). (D) Quantification of the fluorescence intensity of the Syt-1 label. (Mean ± SEM; n = 3 exp., 6 cells per exp.; **p<0.01, paired t-test). Distinct roles of BDNF for NL1 and NL2 induced increases in synapse number and maturation. (E, F) Representative images and quantitation of number of Bassoon puncta per 10 µm dendrite on DIV6 BDNF^lox/lox^ cultures and Cre-recombinase transduced cultures transfected with mOrange, NL1-mOrange (NL1), or NL2-mOrange (NL2) (red) and immunostained for Bassoon (blue). (Mean ± SEM; n = 3 exp., 6 cells per exp.; *p<0.05, **p<0.01, ***p<0.001, paired t-test, significance between conditions was verified with a two-way ANOVA (***p<0.001)). Scale bar is 10 µm. (G, H) Representative images and quantitation of cultured hippocampal neurons from BDNF^lox/lox^ mice with or without transduction with Cre-recombinase (green) transfected with mOrange, NL1-IRES-mOrange (NL1), or NL2-IRES-mOrange (NL2) (red) on DIV4. On DIV6 cultures were stimulated in the presence of antibodies against the lumenal domain of Synaptotagmin-1 (Syt1) to label recycling synaptic vesicles. After washing the cells were fixed and immunolabeled with secondary antibodies against the Syt1-antibody (blue) (mean ± SEM; n = 3 exp., 6 cells per experiment; **p<0.01, ***p<0.001, paired t-test). Scale bar is 10 µm.

To test the functional maturation of presynaptic terminals in DIV15 BDNF depleted cultures, we applied the Syt1-uptake assay. The Syt1-label intensity in Cre transduced cultures was significantly reduced compared to control cultures (Fig. 3C, D), indicating that also functional presynaptic maturation was reduced in BDNF-depleted cultures. Thus, knockout of BDNF impairs presynaptic maturation in a way similar to knockout of NL1.

### NL1 and NL2 fail to induce presynaptic maturation in BDNF knockout cultures

To further corroborate the notion that NL-induced presynaptic maturation requires BDNF, we overexpressed NL1 and NL2 in BDNF depleted mouse cultures on DIV4 and assayed maturation on DIV6. We co-expressed untagged full-length Neuroligins with IRES-driven mOrange, to visualize transfected neurons in GFP-Cre transduced cultures. Applying LatA to mouse cultures significantly reduced the number of Bassoon puncta on mOrange expressing neurons, whereas the number of puncta on NL1 and NL2 overexpressing neurons remained virtually unchanged (Fig. 3E, F), as expected from our results obtained in rat cultures transfected with NL-GFP constructs. Transducing with Cre to remove BDNF had distinct effects on NL1 and NL2 overexpression: BDNF depletion did not affect the number of Bassoon puncta on NL1-overexpressing neurons, while the number of Bassoon puncta on NL2-overexpressing neurons decreased significantly. In contrast, LatA application led to a significant reduction in the number of Bassoon puncta on NL-1 overexpressing neurons and on NL2-overexpressing neurons, indicating that in BDNF-depleted cultures, both Neuroligins failed to induce structural maturation. We then tested synaptic vesicle recycling, in DIV6 BDNF depleted cultures transfected with NL1-mOrange (NL1), NL2-mOrange (NL2), or mOrange on DIV4. Knockout of BDNF significantly decreased the intensity of the Syt1-label in presynaptic terminals compared to mOrange expressing control cells, and virtually prevented the NL-induced increase in SV recycling (Fig. 3G, H).

Is Neurexin binding required for presynaptic maturation? To test this, we overexpressed NL1 and a NL1 variant that does not bind to Neurexins (Ko et al., 2009) in young cultures. Both WT NL1 and the Neurexin binding deficient mutant generated the same Latrunculin resistance and Syt-1 uptake levels when overexpressed at DIV5 in rat cultures (Fig. 4). LatA resistance levels were also similar when we compared the action of both constructs in NL1-KO cultures (Fig. 5).

**Figure 4:**
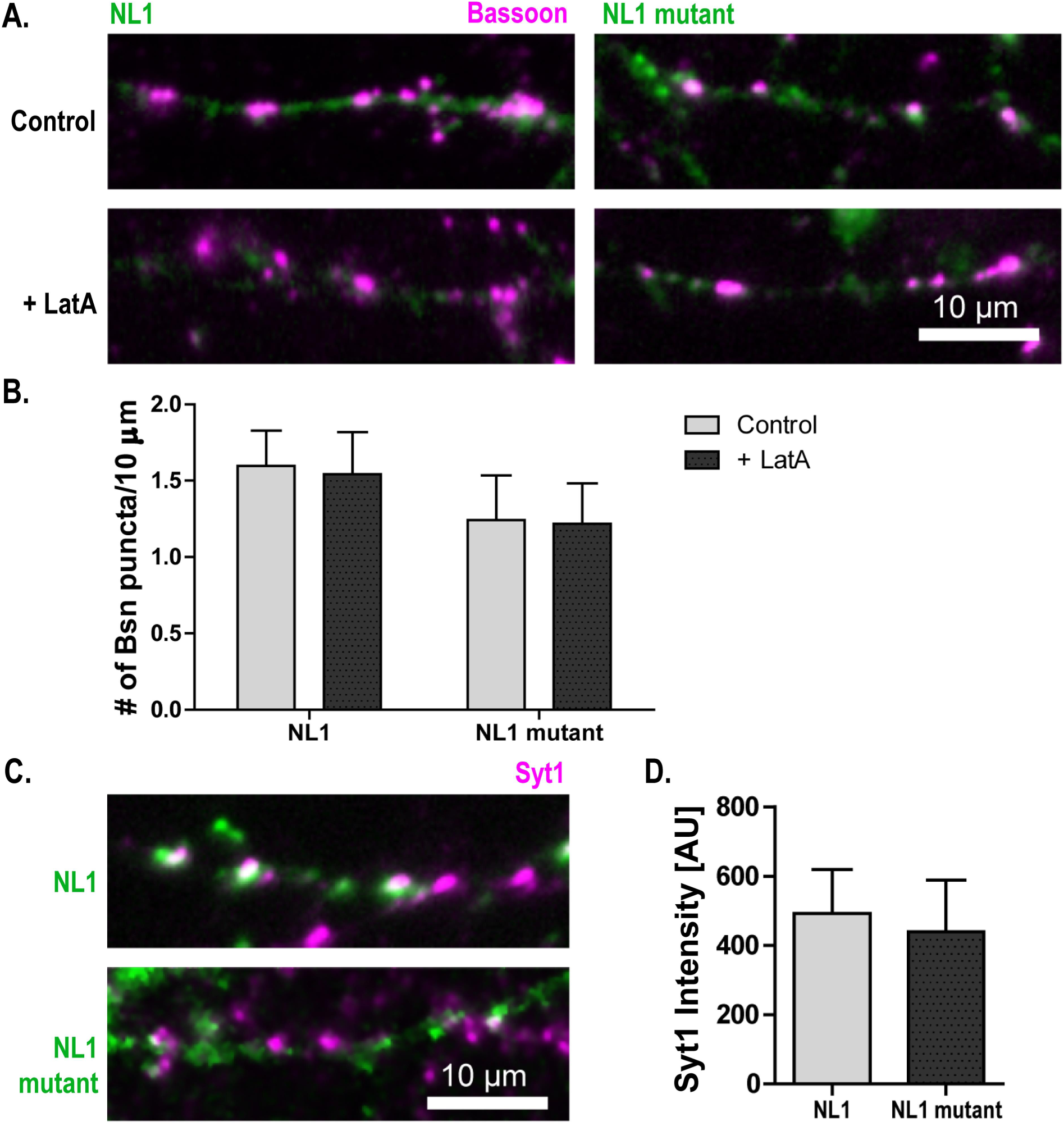
Similar effects of NL1 and a Neurexin binding deficient NL1 mutant on structural and functional presynaptic maturation. (A) DIV5 cultured hippocampal neurons transfected with NL1-IRES-mOrange (NL1) or mutated NL1-IRES-mOrange (NL1 mutant), with and without LatA treatment and immunostained for Bassoon (magenta). (B) Quantification of the number of Bassoon puncta per 10 µm dendrite for the conditions indicated in panel A. (Mean ± SEM; n = 3 exp., 10 cells per exp.; paired t-test). (C) The cultures were stimulated in the presence of antibodies directed against the lumenal domain of Synaptotagmin-1 (Syt1) to label recycling synaptic vesicles (magenta). (D) Quantification of the fluorescence intensity of the Syt-1 label. (Mean ± SEM; n = 3 exp., 10 cells per exp.; paired t-test). Scale bar is 10 µm.

**Figure 5:**
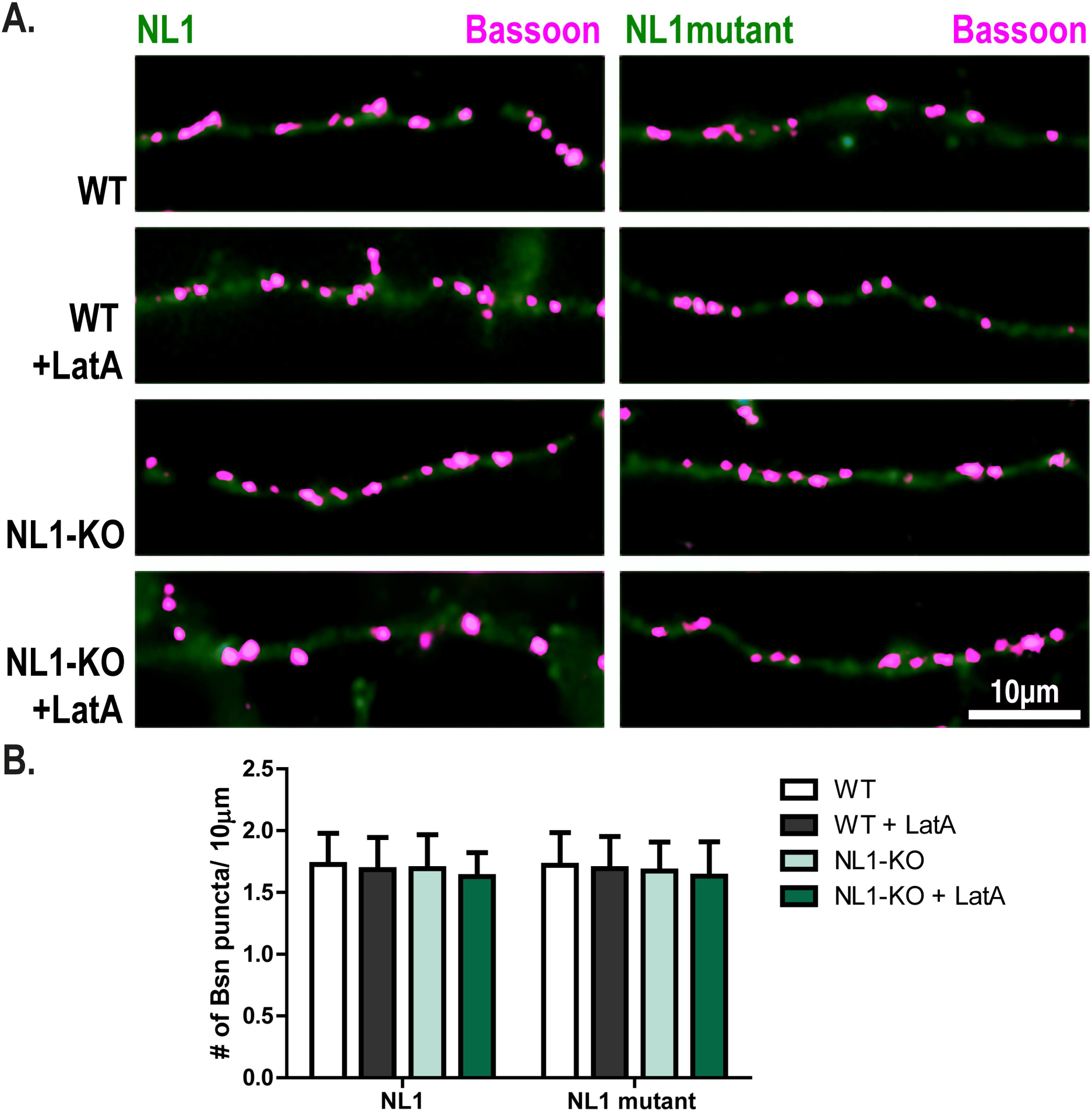
Similar effects of NL1 and a Neurexin binding deficient NL1 mutant on structural presynaptic maturation in NL1-KO cultures. (A) DIV5 cultured cortical WT and NL1-KO neurons transfected with NL1-IRES-mOrange (NL1) or mutated NL1-IRES-mOrange (NL1 mutant), with and without LatA treatment and immunostained for Bassoon (magenta). (B) Quantification of the number of Bassoon puncta per 10 µm dendrite for the conditions indicated in panel A. (Mean ± SEM; n = 3 exp., 10 cells per exp.; paired t-test). Scale bar is 10 µm.

### BDNF rescues defective presynaptic maturation in NL1-KO cultures

Why do NLs fail to promote maturation in the absence of BDNF? One explanation is that NLs elicit or enhance BDNF signaling to promote presynaptic maturation. In this case, exogenous BDNF should rescue maturation in NL1 knockout cultures, which fail to mature (Wittenmayer et al. 2009). To test this, we applied BDNF to advanced cortical cultures (DIV14) from WT and NL1-KO mice and tested them for structural and functional maturation a day later. Remarkably, BDNF application reestablished structural and functional maturation in these NL1-KO cultures within one day (Fig. 6).

**Figure 6:**
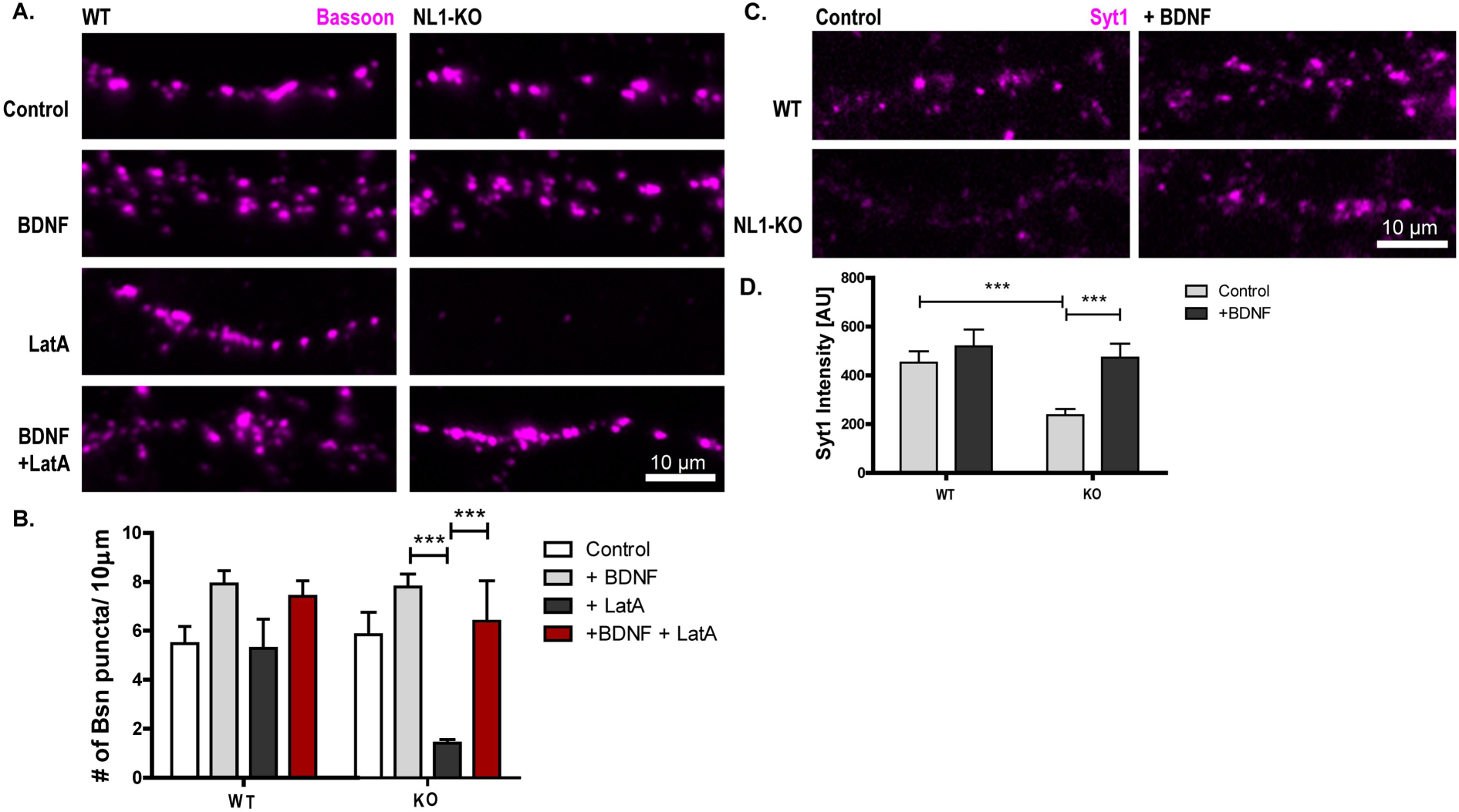
BDNF restores presynaptic maturation in NL1-deficient cultures. (A, B) Representative images and quantification of cultured cortical neurons from WT and NL1-KO mice fixed on DIV15. Prior to fixation, the neurons were treated with either BDNF for 22 hours, LatA for 18 hours, or both (see labeling on the left). Following fixation, the neurons were immunolabeled with a Bassoon antibody (magenta) (mean ± SEM; n = 3 exp., 6 cells per experiment and condition; ***p<0.001, paired t-test). Scale bar is 10 µm. (C, D) Representative images and quantification of cultured cortical neurons from WT and NL1-KO mice fixed on DIV15. On DIV15 cultures were stimulated in the presence of antibodies against the lumenal domain of Synaptotagmin-1 (Syt1) to label the recycling of synaptic vesicles. After washing the neurons were fixed and immunolabeled with secondary antibodies against the Syt-1 antibody (magenta) (mean ± SEM; n = 3 exp., 10 cells per experiment; **p<0.01, ***p<0.001, paired t-test). Scale bar is 10 µm.

### NL1 increases postsynaptic BDNF levels to induce presynaptic maturation

To test whether the BDNF that mediates Neuroligin induced presynaptic maturation is released from the pre- or postsynaptic side, we introduced Cre recombinase and Neuroligins into a small number of neurons from *bdnf* ^lox/lox^ mice by transfection rather than viral transduction. Using this paradigm, only the small number of neurons (dozens per coverslip containing 70000 cells) expressing a Neuroligin also expressed Cre, thus removing BDNF from the postsynaptic cell. To verify co-transfection of NLs and Cre, the NL-constructs co-expressed mOrange and the Cre plasmid co-expressed GFP from an internal ribosomal entry site. As a control, we co-transfected cells with NLs and GFP. We then tested the structural and functional maturation of the boutons contacting these neurons using the LatA- and Syt1-uptake assay (Fig. 7 A–D). At dendrites of NL1 overexpressing neurons co-transfected with Cre, the number of Bassoon stained puncta as well as the Synaptotagmin-1 fluorescence intensity was significantly decreased compared to the neurons co-transfected with GFP as a control, indicating that postsynaptic BDNF mediates NL1-induced presynaptic maturation. In contrast, at dendrites of NL2 overexpressing neurons, we did not observe any change in the number of Bassoon puncta or Synaptotagmin-1 fluorescence intensity, suggesting that in this case postsynaptic BDNF was not needed to mediate presynaptic maturation.

**Figure 7:**
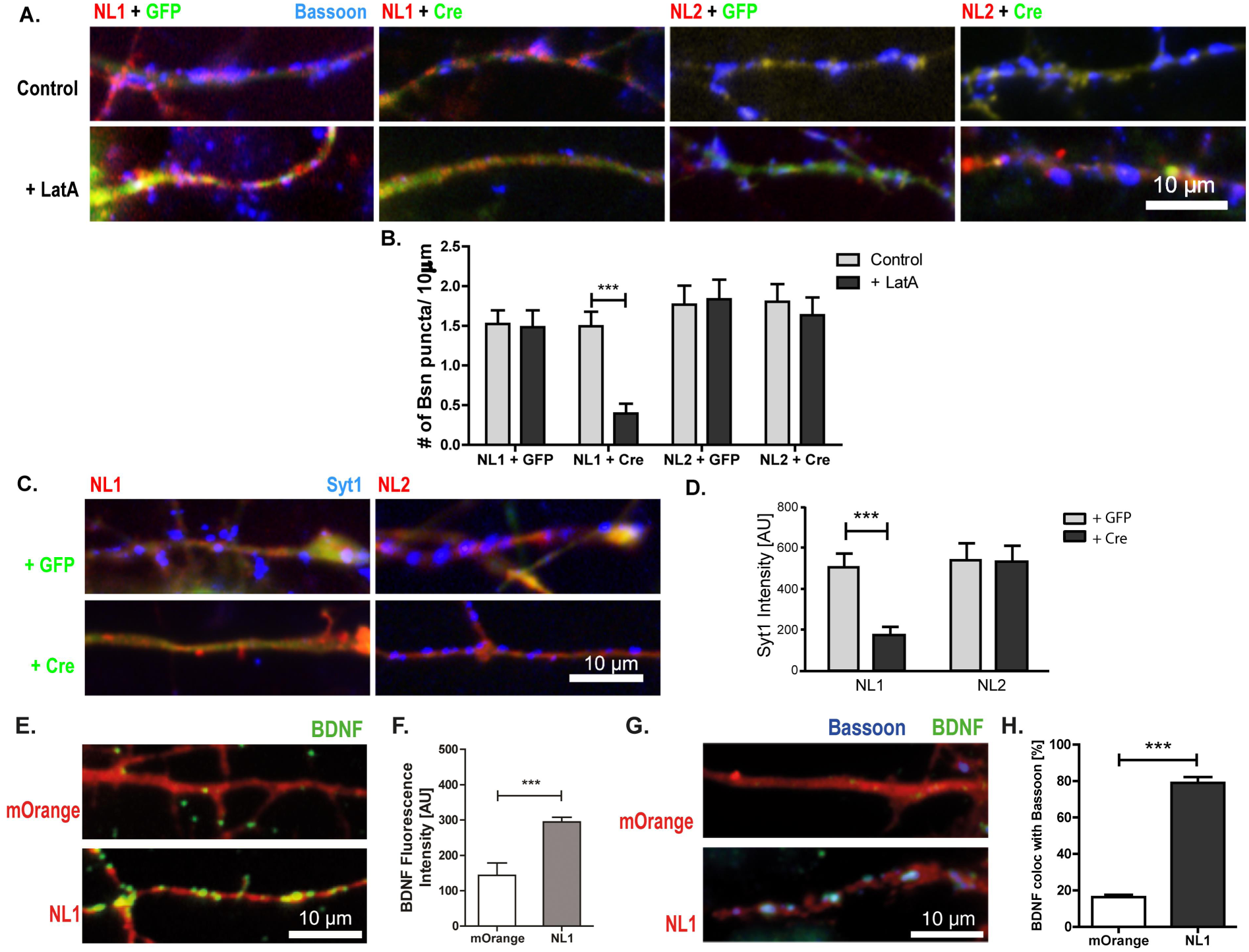
Postsynaptic BDNF is required for NL1-induced, but not for NL2-induced, presynaptic maturation. (A, B) Representative images and quantification of cultured hippocampal neurons from BDNF^lox/lox^ mice co-transfected with either NL1-IRES-mOrange (red) + Cre / GFP (green), or NL2-IRES-mOrange (red) + Cre / GFP (green) on DIV4. Neurons were fixed on DIV6 and immunostained for Bassoon (blue) following a 6-hour LatA treatment to depolymerize F-Actin (mean ± SEM; n = 3 exp., 6 cells per exp.; ***p<0.001, paired t-test). Scale bar is 10 µm. (C, D) Representative images and quantitation of cultured hippocampal neurons from BDNF^lox/lox^ mice co-transfected with either NL1-mOrange (red) + Cre / GFP (green) or NL2-mOrange (red) + Cre / GFP (green) on DIV4. On DIV6 cultures were stimulated in the presence of antibodies against the lumenal domain of Synaptotagmin-1 (Syt1) to label recycling synaptic vesicles. After washing the cells were fixed and immunolabeled with secondary antibodies against the Syt1-antibody (blue) (mean ± SEM; n = 3 exp., 10 cells per experiment; **p<0.01, ***p<0.001, paired t-test). (E, F) Overexpressing NL1 increases BDNF immunofluorescence at active zones. Representative images and quantification of BDNF fluorescence in the dendritic region of neurons expressing mOrange or NL1-IRES-mOrange. On DIV5 the neurons were fixed and immunostained for BDNF (green). The average intensity of fluorescent puncta was quantified. (G, H) BDNF immunosignals colocalize with Bassoon in NL1 overexpressing neurons (mean ± SEM; n = 2 exp., 10 cells per experiment; **p<0.01, ***p<0.001, paired t-test). Scale bar is 10 µm.

Activity-induced expression is a central feature of BDNF, and NL1 can enhance neuronal activity by enhancing NMDA receptor activity. To test whether NL1 leads to an increased expression of BDNF, we transfected DIV3 cultured hippocampal neurons with NL1-IRES-mOrange or mOrange as a control. After fixation, we immunostained the neuronal cultures for BDNF. We observed a significant increase of BDNF fluorescence in the NL1 overexpressing neurons compared to control neurons transfected with mOrange (Fig. 7 E, F), and ~80 percent of these BDNF signals colocalized with Bassoon (Fig. 7G, H).

Together, our data suggest that BDNF is an essential novel participant in NL-mediated presynaptic maturation.

## Discussion

Synapses go through stages of maturation to create fully functional networks. Studying early network formation in cultured neurons, we found that presynaptic maturation requires the concerted action of Neuroligins and BDNF. BDNF application mimicked the boosting effects of Neuroligins on AZ stability and synaptic vesicle recycling. Perturbing endogenous BDNF signaling reduced both ongoing maturation and the enhancing effects of NL1 and NL2. BDNF appeared to act downstream of Neuroligins, permitting exogenous BDNF to restore presynaptic maturation in NL1 knockout neurons. Our data reveal that BDNF and Neuroligins jointly mediate structural and functional presynaptic maturation, thus implicating neurotrophin action in Neuroligin-based signaling.

### Presynaptic maturation: distinct or overlapping pathways?

Three scenarios could account for this cooperative action of NLs and BDNF: first, presynaptic maturation could be accomplished through the combined effects of two independent pathways, one involving NL-based cell adhesion and one involving BDNF signaling, which add up to full maturation. This seems unlikely for functional maturation because the effects of NL1 and NL2 in enhancing SV recycling were virtually abolished when BDNF signaling was blocked. If the pathways were independent, the contribution of NLs to mediating presynaptic maturation should be unaffected. Structural maturation was reduced by 50% in BDNF depleted cultures and by 80% in NL1 deficient cultures at DIV15. If maturation was mediated by two independent pathways, the contributions of BDNF and NLs should not exceed 100%. Of note, we may underestimate the importance of BDNF for structural maturation because the floxed-BDNF cultures were rendered BDNF-depleted via viral delivery of Cre, hence residual BDNF – produced by uninfected neurons or incomplete Cre action – may still exist in the media. In any case, even a cautious estimate, assuming that BDNF is responsible for 50% and NLs for 80% of structural maturation, argues against two independent pathways.

Second, a cell adhesion based pathway and a neurotrophin based pathway each may have the potency to generate full maturation, albeit through seperate molecular cascades. This could reflect a redundant scenario to ensure proper maturation, and the pathways could depend on each other such that one is upregulated when the other one lags behind. We cannot exclude this possibility, but our data show that this potency – if it existed – is not used during synaptogenesis, since the absence of NLs or BDNF in knockouts is not compensated for by the presence of the respective other pathway.

Third, NLs and BDNF may act in a shared pathway or in closely overlapping pathways. Two aspects argue for this scenario: BDNF fully rescued impaired presynaptic maturation caused by the loss of NL1, but NLs did not rescue impaired maturation caused by BDNF deficiency. Thus, NLs and BDNF cannot be acting in seperate pathways, and instead must act in the same or in overlapping pathways. Moreover, overexpression of NL1 increased BDNF expression in the transfected neurons, indicating that the action of NL1 affects BDNF, providing a hint at the site of overlap.

### The overall effects of BDNF and NLs

A joint action of BDNF and NLs appears to be plausible both based on our new observations and on the known actions of these molecules. BDNF application increased the number of Bassoon puncta and AZ stability in young cultures, and BDNF deprivation prevented the acquisition of the mature state in advanced cultures. These effects resemble the effects of NL1 overexpression and NL1 knockout, respectively (Wittenmayer et al. 2009), arguing for the two molecules acting together during synapse maturation. Overexpressing NL2 had virtually identical potency suggesting that accelerating presynaptic maturation is a general property of Neuroligins.

Several known properties of BDNF are reminiscent of the effects of NLs. For example, BDNF and NL1 increase the levels of SV proteins (Wittenmayer et al. 2009; Hoy et al. 2013; Pozzo-Miller et al. 1999; Tartaglia et al. 2001). BDNF and NL1 also increase release probability (William J Tyler et al. 2006; Wittenmayer et al. 2009). These similarities between the proteins inspired us to study whether BDNF and NLs act in concert to mediate presynaptic maturation.

BDNF increased the number of Bassoon puncta in our young cultures. This is consistent with increased synapse numbers in advanced (DIV14) cultures and postnatal (P7) slice cultures treated with BDNF (Vicario-Abejón et al. 1998; W J Tyler and Pozzo-Miller 2001). We found that both NL1 and NL2 needed BDNF signaling to induce presynaptic maturation, but only NL2 also needed BDNF to increase the number of synapses. This is consistent with the particular importance of NL2 and BDNF for inhibitory synapse formation or maintenance (Chih, Engelman, and Scheiffele 2005; Chubykin et al. 2007; Aguado et al. 2003; Kohara et al. 2007). Interestingly, BDNF was not required for the increase in synapse number induced by NL1 overexpression, indicating that NL1 and NL2 use distinct pathways to increase the number of synapses, but similar, BDNF-dependent, ways to increase maturation.

The importance of BDNF for both Neuroligin-induced and endogenous maturation of SV recycling is consistent with the known role of BDNF in increasing the number of docked SVs and the extent of SV recycling in advanced stages (≥ DIV 13) of culture development (Collin et al. 2001; Shinoda et al. 2014) and in P7 slice cultures (Tartaglia et al. 2001). These effects of BDNF are usually interpreted as a modulatory element acting in existing networks. Our data indicate that this potency of BDNF is also a crucial component of NL-mediated presynaptic maturation.

BDNF also mimicked and mediated – at least in part – the effects of NL1 and NL2 on AZ stabilization after F-actin disruption by LatA. In addition, knockout of either BDNF or NL1 prevented the acquisition of the structurally mature state, as shown before for NL1 (Wittenmayer et al. 2009). Overall, our data show that BDNF is a necessary component for the NL-mediated effects on structural and functional maturation. It is unlikely that BDNF acts as a generally permissive element, for example by promoting axonal outgrowth and dendritic arborization, because the number of AZs was not reduced in advanced BDNF depleted cultures: like in NL1 KO cultures, presynaptic terminals formed and decorated dendrites at a normal density, but remained at an immature state. We propose that NLs harness the positive effects of BDNF signaling on presynaptic terminals to boost presynaptic maturation during early stages of network formation.

### How do NLs and BDNF interact?

We have previously shown that structural maturation requires both NL1 and NMDAR activity, suggesting that neuronal activity constitutes an additional element in the process of synaptic maturation (Wittenmayer et al. 2009). Notably, both NL1 and BDNF recruit NMDA receptors (NMDARs) to synapses (Chih et al. 2005; Chubykin et al. 2007; Caldeira et al. 2007; Wittenmayer et al. 2009; Budreck et al. 2013). In our study, overexpression of NL1 increased the fluorescence intensity of BDNF immunosignals. NL1 might increase network activity by recruiting NMDARs, thereby allowing for activity-dependent BDNF expression and secretion. Subsequently, BDNF could further enhance NMDAR activity, Ca^2+^ influx and downstream signaling. Ca^2+^ signaling, via CaMKII, promotes NL1 surface expression and function, as well as BDNF expression and secretion (Bemben et al. 2014; Zhou et al. 2006; Kolarow, Brigadski, and Lessmann 2007). Thus, NMDAR / CaMKII signaling could link NL1 and BDNF action in presynaptic maturation.

In which sequence do NL1 and BDNF act? While BDNF application rescued NL1 deficiency, NL1 overexpression failed to rescue BDNF deficiency. This indicates that BDNF is downstream of NL1 action. This is consistent with the upregulation of BDNF by NL1. Intriguingly, since neuronal activity is an element in the process, a circular system could arise, where BDNF and NL1 stimulate each other’s actions. In particular, NL1 could enhance NMDA receptor-dependent neuronal activity and CaMKII signaling, thus increasing both BDNF expression and secretion. This in turn increases surface expression of NMDAR, thus closing the feedback loop. In addition, the positive regulation of NL1 via CaMKII (Bemben et al. 2014) further strengthens the loop.

In this scenario, NL1 is the only protein that is exclusively postsynaptic. Thus, such a loop could involve both sides of the synapse, but is likely triggered postsynaptically by NL1. An interesting finding for us was that excision of BDNF in a small number of postsynaptic neurons had contrasting effects on maturation induced by NL1 or NL2, respectively. Whether BDNF is released from axons or dendrites has been controversial because secretion from either side of the synapse has been observed. We found that presynaptic inputs induced by NL1 overexpression in BDNF depleted neurons failed to mature, while BDNF was not required in neurons overexpressing NL2. We do not know an obvious reason for this difference, but it may relate to the different modes observed for excitatory versus inhibitory synapse assembly (Wierenga, Becker, and Bonhoeffer 2008) or the distinct postsynaptic partners that NL1 and NL2 recruit (Krueger et al. 2012). Our data show that NL2 does not rely on BDNF in the transfected neuron, indicating that either global BDNF levels or presynaptic BDNF are sufficient for NL2 mediated maturation. In contrast, NL1 action exclusively relied on locally produced BDNF released from the transfected neuron. While BDNF seems to be exclusively presynaptic in the hippocampal CA3 and CA1 of 8 week old animals (Dieni et al. 2012), our findings for NL1-dependent maturation are consistent with dendritic release in cultured neurons (Hartmann, Heumann, and Lessmann 2001; Kuczewski et al. 2008) and – in the context of long-term synaptic plasticity – in intact brain tissue (Edelmann et al. 2015; Hedrick et al. 2016; Harward et al. 2016). Overall, our data indicate that NL1 induces presynaptic maturation via dendritic release of BDNF.

### Potential presynaptic mechanisms

What are the presynaptic events that follow? Notably, some actions of Neuroligins are dependent on presynaptic Neurexins, while others are not (Ko et al. 2009): A mutated NL1 that cannot bind to Neurexins failed to increase synapse size when expressed in cultured neurons while still able to increase synapse number and strength. We found that WT NL1 and the Neurexin binding deficient mutant showed the same potency for inducing presynaptic maturation, both with respect to LatA-resistance and SV recycling. Thus, Neurexin binding may be dispensible for NL1/BDNF-induced presynaptic maturation. In contrast, binding of BDNF to TrkB receptors is likely involved because scavenging BDNF with soluble TrkB-Fc bodies impaired maturation. Interestingly, BDNF treatment increases F-actin polymerization via TrkB/LIMK1 interaction during axonal outgrowth (Dong et al. 2012), raising the possibility that acquisition of LatA-resistance could be due to enhanced stabilization of F-actin.

That the architecture of synapses becomes independent of F-actin during development was originally found by Zhang and Benson (Zhang and Benson 2001), and later shown to involve N-Cadherin signaling (Bozdagi et al. 2004). This is remarkable, since N-Cadherin recruits NL1 to synapses (Stan et al. 2010), suggesting that N-Cadherin/NL1-based cell adhesion triggers BDNF-signaling to induce LatA-resistance of AZs. The biological purpose of LatA-resistance is unclear. Also, what replaces F-actin as a stabilizing compound during neuronal development is unclear. At least two principal mechanisms are conceivable: a) the AZ could become stabilized by transsynaptic anchoring involving direct physical connections to the PSD; b) AZ molecules could become interconnected, thus tightening the cytomatrix of the active zone inherently. Impairment of either of these mechanisms could account for the reduced LatA-resistance as assayed in BDNF and NL1 KO cultures. For our gain-of-function experiments a third option cannot be excluded: application of BDNF and overexpression of NLs may increase the levels of F-actin thus enhancing F-actin stabilization so that some of it survives the LatA-treatment. Synaptic sites of extremely stable F-actin have been previously observed (Zhang and Benson 2001).

In either case, i.e. through increased F-actin levels or through increased inherent stability, NLs and BDNF could make AZs independent of ongoing actin remodeling and thus allow for a simple mechanism to stabilize mature AZs while remodeling mechanisms continue. Conversely, before LatA-resistance is established, i.e. in the structurally immature state, AZs may be particularly malleable. In cultures from NL1 KO mice, which have normal synapse numbers but fail to acquire LatA-resistance, the tenacity of excitatory synapses is reduced. The turnover of synaptic proteins and the fluctuation of their concentration at synapses is increased, and synaptic activity destabilizes synapses in NL1 KO cultures (Zeidan and Ziv 2012). The LatA-resistant state may thus protect synapses against activity-induced changes in mature cultures, and the LatA-sensitive state may allow for remodeling in early cultures.

### Outlook

Activity-dependent synapse maturation is a central event in network refinement, and failures in this process likely contribute to neurodevelopmental disorders such as ASD. Interestingly, many proteins implicated in ASD are regulated by neuronal activity, emphasizing the importance of understanding the underlying signaling pathways and identifying convergent pathways. Both NL and BDNF are regulated in an activity-dependent manner and initiate processes that further enhance neuronal/network activity (Ebert and Greenberg 2013). Through their interaction in presynaptic maturation a new site of convergence between two signaling systems emerges. This novel insight has concrete benefits: defects in both NL function and BDNF secretion have been implicated in ASD in man and behavioral abnormalities in mice (Südhof 2008; Sadakata et al. 2007; Sadakata et al. 2012). Thus, the potency of BDNF to overcome the lack of maturation caused by NL1-deficiency provides an opportunity for therapeutic strategies. This interplay might extend beyond synapse maturation because Neuroligin-1 also promotes survival of new-born neurons in the adult hippocampus (Schnell et al. 2014).

At the cell biological level, our data suggest a scenario where Neuroligins interact with Neurexins to promote synapse formation, and subsequently switch to a different receptor or even act without a presynaptic receptor to induce BDNF release and synaptic maturation. The multiple roles of BDNF for brain development and plasticity require tight spatial and temporal control of BDNF expression and release. Neuroligins could provide this control in the context of presynaptic maturation, by linking cell adhesion and synaptic activity to local BDNF secretion.

It will be interesting to see whether this transsynaptic teamwork between Neuroligins and BDNF is also effective during synapse maturation in the adult brain, e.g. during constitutive synapse turnover, plasticity related synapse formation, and adult neurogenesis.

## Acknowledgements

Lentivirus production was kindly performed by Tania Meßerschmidt. We thank Martin Rothkegel (TU Braunschweig) for his help in the construction of the virus, as well as Dilja Krueger and Liam Tuffy (Max-Planck-Institute for Experimental Medicine, Göttingen) for providing NL1-KO mice, and Irmgard Weiß for excellent technical assistance. This work was supported by the DFG via the DFG Research Center for Nanoscale Microscopy and Molecular Physiology of the Brain (CNMPB) to T.D, and via KO 1674/5–1 to M.K.

## Author Contributions

Conceptualization, T.D.; Methodology, T.D. and A.P.; Formal analysis, A.P.; Investigation, A.P. and N.G.; Resources, T.D., M.K. and N.G.; Writing – Original Draft, T.D. and A.P.; Writing – Review & Editing, T.D., M.K., A.P. and N.G.; Visualization, A.P.; Supervision, T.D.; Project Administration, T.D. and A.P.; Funding Acquisition, T.D. and M.K.

## Competing interests

The authors declare no competing interests.

